# High-resolution estimates of crossover and noncrossover recombination from a captive baboon colony

**DOI:** 10.1101/2021.07.10.451667

**Authors:** Jeffrey D. Wall, Jacqueline A. Robinson, Laura A. Cox

## Abstract

Homologous recombination has been extensively studied in humans and a handful of model organisms. Much less is known about recombination in other species, including non-human primates. Here we present a study of crossovers and non-crossover (NCO) recombination in olive baboons (*Papio anubis*) from two pedigrees containing a total of 20 paternal and 17 maternal meioses, and compare these results to linkage-disequlibrium (LD) based recombination estimates from 36 unrelated olive baboons. We demonstrate how crossovers, combined with LD-based recombination estimates, can be used to identify genome assembly errors. We also quantify sex-specific differences in recombination rates, including elevated male crossover and reduced female crossover rates near telomeres. Finally, we add to the increasing body of evidence suggesting that while most NCO recombination tracts in mammals are short (e.g., < 500 bp), there are a non-negligible fraction of longer (e.g., > 1 Kb) NCO tracts. We fit a mixture-of-two-geometric distributions model to the NCO tract length distribution and estimate that >99% of all NCO tracts are very short (mean 24 bp), but the remaining tracts can be quite long (mean 11 Kb). A single geometric distribution model for NCO tract lengths is incompatible with the data, suggesting that LD-based methods for estimating NCO recombination rates that make this assumption may need to be modified.

**Significance:** Most homologous recombination events are noncrossovers (NCO), but little is known about NCO conversion tract lengths. Here we utilize whole-genome sequence data from large baboon pedigrees to estimate the NCO tract length distribution and to study other aspects of recombination.

## Introduction

Homologous recombination is a fundamental biological process, thought to be necessary for the proper segregation of chromosomes during meiosis and essential for the efficacy of natural selection. Recombination rates in higher eukaryotes are generally measured using (1) genetic comparisons between parents and offspring (e.g., using genotype or sequence data), (2) genotyping or sequencing of single or pooled sperm (i.e., potential gametes), or (3) indirect estimation via statistical methods that quantify the relationship between linkage disequilibrium and recombination. Each of these three approaches involve tradeoffs regarding cost/effort and the breadth and depth of information they can provide. In particular, only pedigree-based studies provide both sex-specific recombination estimates and direct identification of both crossover (CO) and non-crossover (NCO) recombination events, but they are more difficult to conduct due to sample acquisition challenges.

Recombination is thought to arise from double strand breaks (DSB) that occur after the pairing of homologous chromosomes during meiosis. Depending on how these breaks are resolved, the result can either be CO recombination, which involves the reciprocal transfer of large chromosomal regions between homologs, and NCO recombination (colloquially called ‘gene conversion’), involving the non-reciprocal replacement of short tracts of DNA from one homolog to another (Orr-Weaver et al. 1981; Szostak et al. 1983). Since crossovers are also associated with gene conversion tracts at the DSB location, we will use the term NCO recombination to describe homologous gene conversion not associated with a nearby crossover.

Theory predicts a close relationship between recombination and patterns of linkage disequilibrium (LD), since homologous recombination will tend to shuffle haplotypes and break down allelic associations. Population genetic analyses of dense genotype and sequence data, along with sperm typing studies, have shown that most human crossovers happen in narrow (1-2 Kbp) ‘hotspots’ (e.g., Chakravarti et al. 1984; Jeffreys et al. 2001; Crawford et al. 2004; Myers et al. 2005), and that this fine-scale structuring of recombination rates can help explain the block-like structure of LD in many parts of the genome (Wall and Pritchard 2003). In most vertebrates, the locations of these hotspots are mediated by the zinc finger *PRDM9* (reviewed in Paigen and Petkov 2018), and recombination hotspot locations are generally not shared across closely related species (e.g., Ptak et al. 2005; Auton et al. 2012; Stevison et al. 2016).

Much less is known about NCO recombination. A handful of studies in humans and model organisms have found that most recombination events are NCOs, but mean tract lengths are quite short – tens or hundreds of base pairs (e.g., Jeffreys and May 2004; Baudat and de Massy 2007; Cole et al 2010; Comeron et al. 2012; Wijnker et al. 2013; Li et al. 2019). This short tract length makes NCO recombination especially difficult to study. In particular, for species with low levels of heterozygosity (e.g., most mammals) many NCO tracts are undetectable because the donor and converted sequences are identical. In most of the remainder only a single heterozygous site is converted, making NCO recombination difficult to distinguish from simple genotype/sequencing errors. While statistical methods have been developed for estimating NCO recombination parameters indirectly from segregating patterns of genetic variation (e.g., Frisse et al. 2001; Gay et al. 2007; Yin et al. 2009; Padhukasahasram and Rannala 2013), these methods are not very accurate primarily because of the small/negligible effect that most NCO tracts have on patterns of genetic variation. In addition, these methods generally assume that NCO tract lengths follow a geometric distribution, which may not be biologically realistic. Because of this, studies of NCO recombination have generally focused on identifying events by comparing the patterns of genetic inheritance of offspring (or potential offspring in the case of sperm typing) from their parents (e.g., Jeffreys and May 2004; Comeron et al. 2012; Wijnker et al. 2013; Williams et al. 2015; Halldorsson et al. 2016; Li et al. 2019).

Among mammals, NCO recombination has been most-studied in humans, with several sperm typing studies (Jeffreys and May 2004; Jeffreys and Neumann 2005; Webb et al. 2008; Odenthal-Hesse et al. 2014), two large pedigree-based studies (Williams et al. 2015; Halldorsson et al. 2016), and a study of genetic variation in autozygous tracts of consanguineous individuals (Narasimhan et al. 2017). Two observations from these studies stand out. First, both pedigree-based studies found evidence for complex NCO events, involving multiple non-contiguous gene conversion tracts that are physically near each other, from the same meiosis, and not associated with a nearby CO (Williams et al. 2015; Halldorsson et al. 2016). Second, both studies also found evidence for apparent long (i.e., > 20 Kbp), contiguous NCO tracts. If real, these long tracts are suggestive of a separate molecular mechanism distinct from the gene conversion expected under the standard DSB model. It is possible though that they reflect a rare, CO interference independent recombination process, or that they are actually complex NCO events with smaller tract sizes that are miscalled due to low marker density.

In this study, we examine patterns of recombination with a focus on NCO tracts using olive baboons (*Papio anubis*) in the baboon colony housed at the Southwest National Primate Research Center (SNPRC). We generate and analyze high-coverage whole-genome sequence data from two pedigrees with large sib-ships (Figure 1), which allows us to estimate sex-specific recombination rates, identify NCO recombination events, and evaluate the long-range accuracy of the current Panubis1.0 genome assembly (cf. Batra et al. 2020). This assembly used Hi-C contact data to join contigs into scaffolds, and the low-resolution linkage map we generate here allows us to assess the accuracy of this approach.

**Figure 1.**
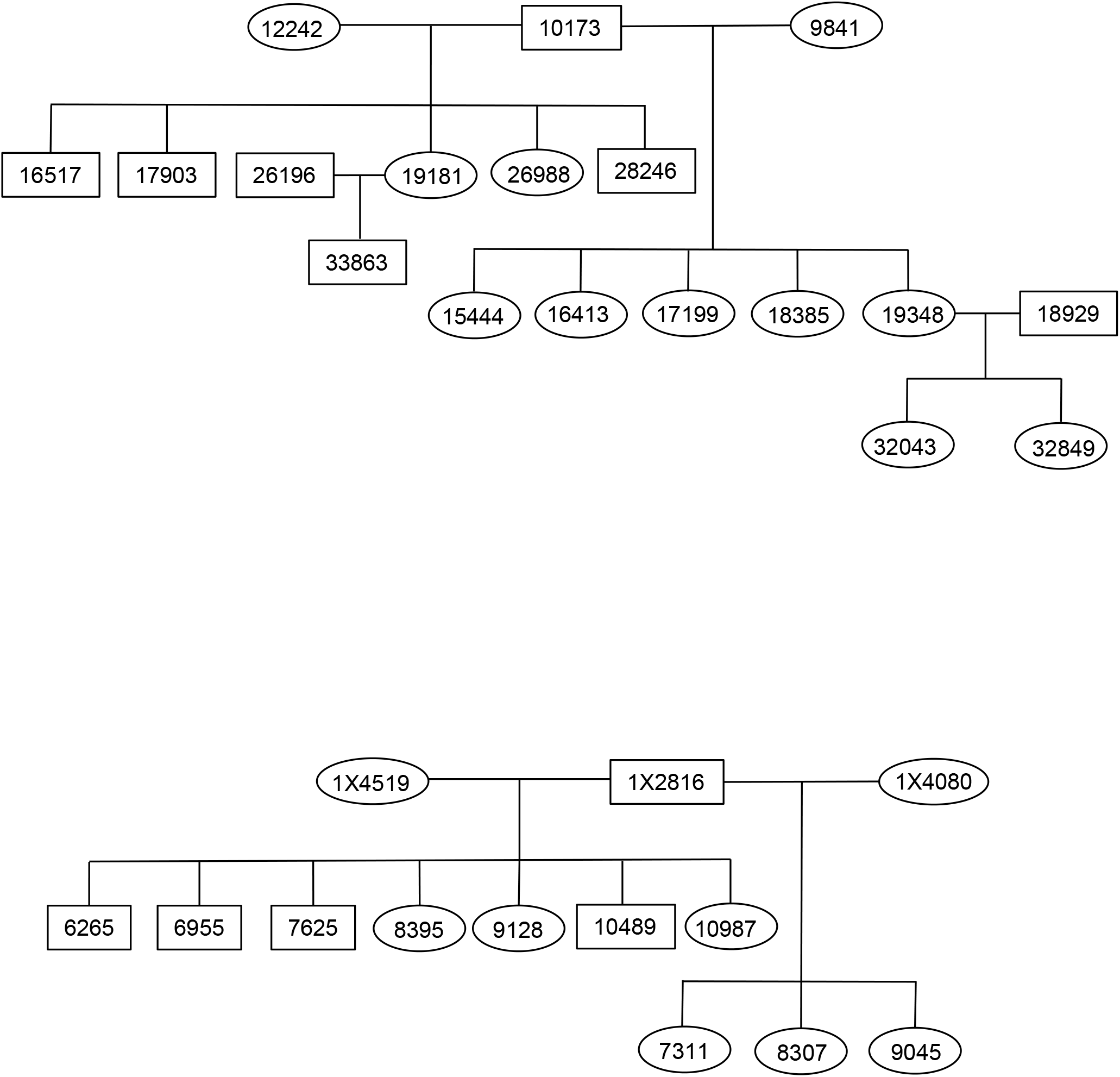
Schematics of the two baboon pedigrees used in this study.

Our choice of baboons was motivated in part by the availability of an extremely large pedigreed colony at the SNPRC, as well as the higher levels of diversity found in baboons relative to humans (e.g., Robinson et al. 2019). Our expectation is that the increased marker density will provide greater resolution on the size distribution of NCO tracts, and that our study of a nonhuman primate will help elucidate whether some of the specific recombination patterns observed in humans can be generalized to a wider group of species.

## Results

### Baboon genetic map

We identified crossovers and NCO recombination events from a total of 20 paternal and 17 maternal meiosis (Supplementary Table S1). In total, we identified 842 autosomal crossovers with a median resolution (i.e., the size of the region over which the crossover location could be placed) of 7.7 Kb. This corresponds to a sex-averaged autosomal genetic map length of 2,293 cM (2,080 cM in males, 2,506 cM in females). Our estimate was 16% larger than a previous estimate based on microsatellite data (Rogers et al. 2000), which reflects both the longer and more complete baboon genome assembly that we used and the much greater marker density of our study. Overall, our results are consistent with the growing body of evidence suggesting that old world monkeys have shorter genetic map lengths, as measured by direct identification of crossovers in pedigrees, than do humans and great apes (e.g., Broman et al. 1998; Rogers et al. 2000, 2006; Kong et al. 2002; Jasinska et al. 2007; Venn et al. 2014).

We also estimated local recombination rates from patterns of LD in 36 unrelated olive baboons using pyrho (Spence and Song 2019). We found that rate estimates are significantly higher within distal regions (≤10 Mb from chromosome ends) relative to proximal regions (>10 Mb from chromosome ends) (two-sided Mann-Whitney U test, *p* < 2.2 * 10^-16^, Supplementary Figure 1).

### Identifying potential genome assembly errors

In performing quality control for our genetic map, we identified several abnormal apparent crossover patterns that likely reflect errors in the Panubis1.0 genome assembly (Figure 2). These included a total of 16 potential inversions, 3 misplaced contigs and 1 potential translocation (Supplementary Table S2). We then used LD-based estimates of recombination using pyrho (Spence and Song 2019) to examine whether patterns of LD provided any additional support. On average, estimated recombination rates at putative synteny breaks are roughly 20 times higher than the estimated rates in the flanking sequences (Figure 3A), consistent with the decrease in LD expected across genome assembly error breakpoints. For 6 proposed inversions and the translocation (Supplementary Table S2), pyrho estimates provide corroborating evidence in finding low levels of estimated recombination (i.e., evidence for synteny) across the ‘corrected’ breakpoints (Figure 3B).

**Figure 2.**
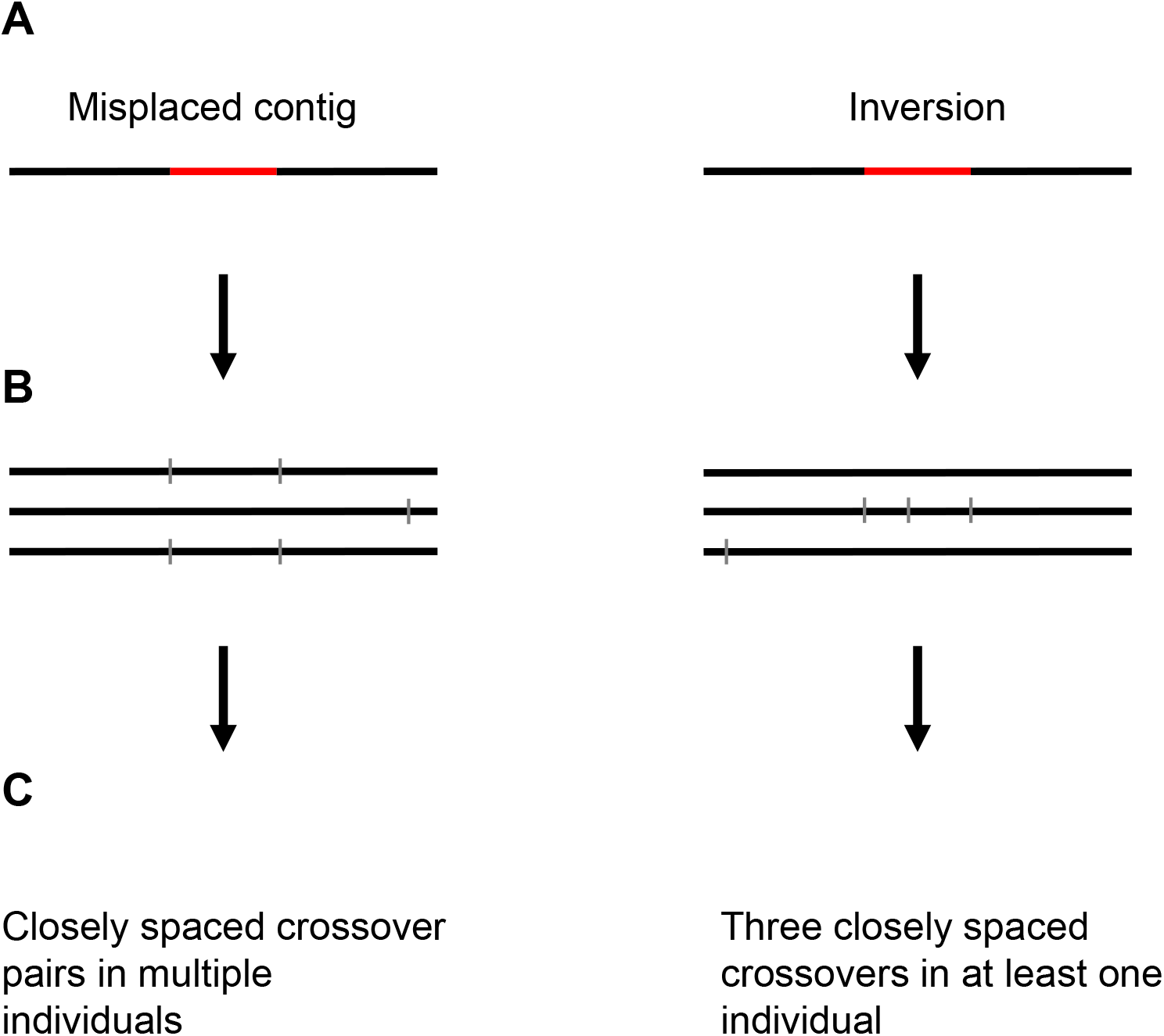
Detecting assembly errors from abnormal crossover patterns. (A) Type of assembly error, (B) Pattern of inferred crossovers in offspring, and (C) Description of pattern.

**Figure 3.**
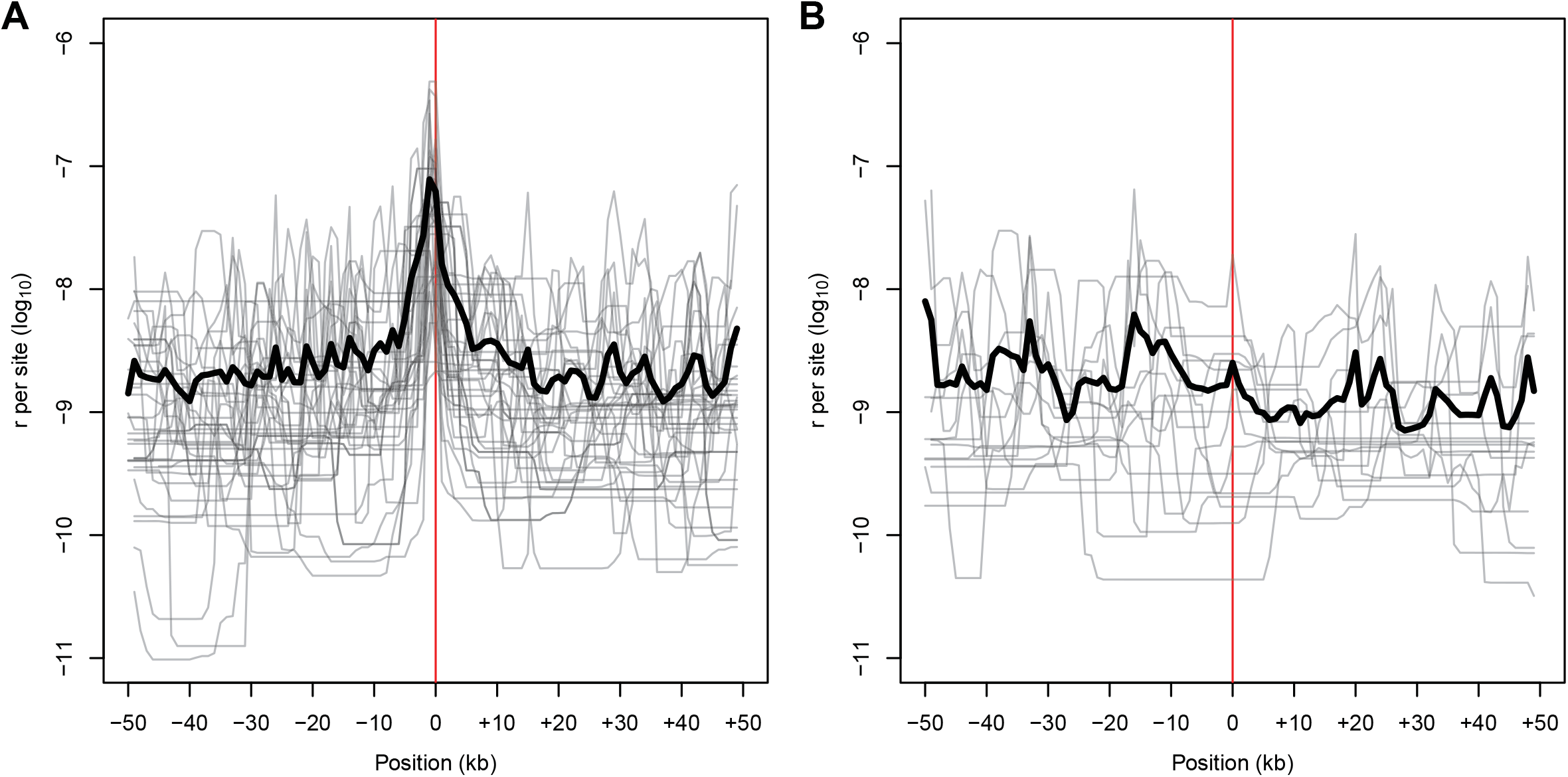
Estimates of ρ from patterns of LD at (A) Breaks of synteny in the current baboon assembly identified from abnormal crossover patterns, and (B) Regions where the Panubis1.0 assembly has been “corrected”

### NCO recombination

After stringent filtering, we identified a total of 325 sites contained in 263 tracts (Supplementary Table S3) that were inferred to be converted due to NCO recombination in tracts < 10 Kb in length. Of the 39 events involving the conversion of more than one heterozygote, the minimal length of the inferred NCO tract was generally small (median = 42 bp), but had a long tail of occasionally longer tracts (mean = 167 bp, including 10 tracts longer than 1 Kbp).

Overall, we estimated a sex-averaged NCO rate of 7.52 * 10^-6^ per site per generation (paternal NCO rate = 5.34 * 10^-6^ and maternal NCO rate = 9.71 * 10^-6^). As with previous human studies (Williams et al. 2015; Halldorsson et al. 2016), we found a handful of more complex NCO recombination events, including 7 regions containing multiple non-contiguous NCO tracts and 9 NCO regions associated with a nearby crossover (Supplementary Table S3; note that 3 regions involve non-contiguous NCO tracts that are also associated with a nearby crossover). In addition, we identified 10 regions consistent with a potential NCO tract of length 10 – 100 Kb (Table 1). Of these, 6 were identified as potential inversion errors in the underlying genome assembly, and three others overlapped with non-inversion potential genome assembly errors (Supplementary Table S2). If we include the remaining long NCO tract into the rate calculation, the estimated sex-averaged NCO rate increases to 8.01 * 10^-6^ per site per generation (paternal NCO rate = 5.34 * 10^-6^ and maternal NCO rate = 1.07 * 10^-5^). These estimates are roughly comparable to NCO rate estimates in humans (e.g., Williams et al. 2015; Halldorsson et al. 2016; Narasimhan et al. 2017).

**Table 1.** List of apparent NCO tracts longer than 10 Kbp

| Chrom. | Parent | Offspring | Minimum tract length | # of NCO sites | Overlap with inversion or BOS? |
| --- | --- | --- | --- | --- | --- |
| 3 | 9841 | 19348 | 40934 | 22 | No |
| 4 | 9841 | 15444 | 24809 | 39 | Yes |
| 6 | 10173 | 18385 | 55304 | 62 | Yes |
| 7 | 10173 | 15444 | 39253 | 2 | Yes |
| 7 | 12242 | 26988 | 45787 | 22 | Yes |
| 8 | 12242 | 28246 | 61766 | 9 | Yes |
| 11 | 1X2816 | 10489 | 24481 | 4 | Yes |
| 13 | 1X2816 | 8307 | 84614 | 54 | Yes |
| 13 | 10173 | 16517 | 86445 | 48 | Yes |
| 16 | 12242 | 17903 | 12403 | 19 | Yes |

### GC bias of NCO tracts

GC-biased gene conversion (gBGC) is a selectively neutral process whereby gene conversion events containing an AT/GC heterozygote in the parent are preferentially resolved to contain the G or C allele in the gamete (Galtier and Duret 2007; Duret and Galtier 2009). Both sperm typing studies (Odenthal-Hesse et al. 2014) and pedigree-based studies (Williams et al. 2015; Halldorsson et al. 2016) in humans have quantified the strength of gBGC in humans. Of the 224 NCO tracts that were informative on gBGC in our study, 129 of them (57.6%) show a transmission bias toward G or C alleles. While this proportion is significantly more than 50% (*p* = 0.014, one-tailed binomial test), it is also significantly less than (*p* = 6.8 * 10^-4^, one-tailed binomial test) the 68% GC bias estimated from human pedigree studies (Williams et al. 2015; Halldorsson et al. 2016).

### Age vs. recombination rate

Previous human recombination studies have documented increases in both CO rate (Kong et al. 2004; Martin et al. 2015) and NCO rate (Halldorsson et al. 2016) with increasing maternal age. While we are underpowered to detect any true correlations between recombination rate and parental age, we did find a marginally significant association between NCO rate and paternal age (*p* = .036; raw data in Supplementary Table S3). All other comparisons of CO or NCO rate with paternal or maternal age were not significant (*p* > 0.1).

### Regional variation in recombination rates

We tabulated the relative numbers of CO and NCO recombination events as a function of distance from telomeres, and stratified the results by sex. We then compared these with sex-averaged recombination rate estimates based on patterns of linkage disequilibrium (Figure 4). As with humans, we find that the male / female CO ratio is higher in distal regions and lower in proximal regions further from the chromosome ends. Near baboon telomeres, males have significantly higher CO rates *and* females have significantly lower CO rates (Figure 4A). We observe a significantly higher male NCO rate near the ends of chromosomes as well (Figure 4B), but did not observe any correlation between female NCO rate and chromosome ends or between CO rate and NCO rate. Consistent with a (partial) decoupling of local CO and NCO rates, we find that pyrho recombination rate estimates are higher near inferred CO locations (Figure 5A) than near inferred NCO locations (Figure 5B).

**Figure 4.**
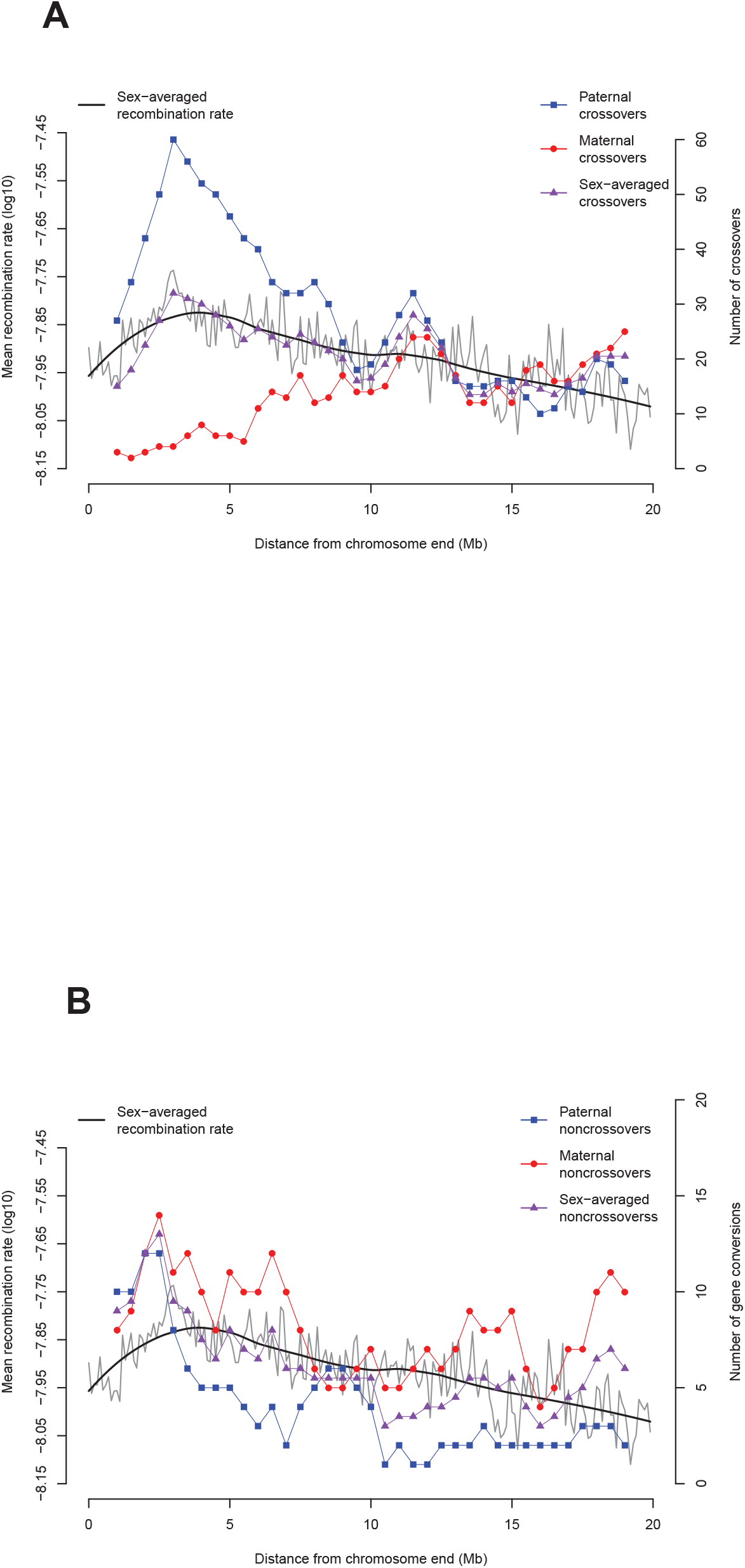
Recombination rates as a function of distance from telomeres. Comparison of LD-based recombination estimates (grey and black) with paternal (blue) and maternal (red) (A) crossover counts, and (B) NCO recombination counts.

**Figure 5.**
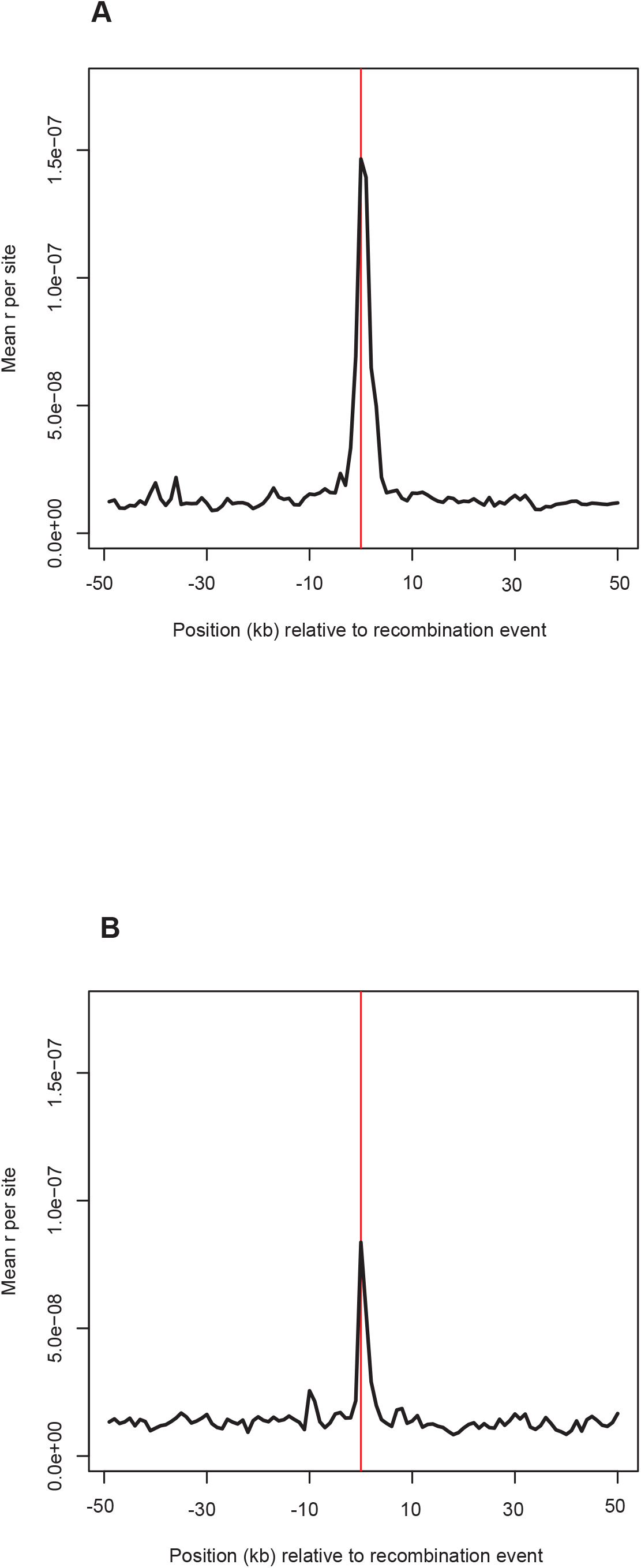
Elevation of pyrho recombination rate estimates near the sites of (A) crossover and (B) NCO recombination events.

### NCO tract length distribution

We used a maximum-likelihood approach for estimating the NCO tract length distribution from the observed patterns of converted NCO sites. We first assumed a geometric tract length distribution, similar to previous studies (e.g., Hilliker et al. 1994; Gay et al. 2007; Miller et al. 2012; Li et al. 2019). If we confine our analyses to NCO tracts less than 10 Kbp in length, we estimate a mean tract length of 309 bp (95% CI = 290 – 341 bp). However, we found that our estimate was roughly proportional to the minimum length of the longest NCO tract in our data set. For example, if we only consider NCO tracts < 5 Kbp in length the estimate is 182 bp (95% CI = 173 – 201 bp), or for tracts < 1 Kbp the mean length estimate is 58 bp (95% CI = 48 – 71 bp). This is in part because the geometric distribution does not fit the data well. In particular, while most NCO sites are consistent with a short (i.e., < 100 bp) tract length, there is a tail of longer NCO tracts that must be kilobases long (Figure 6A).

**Figure 6.**
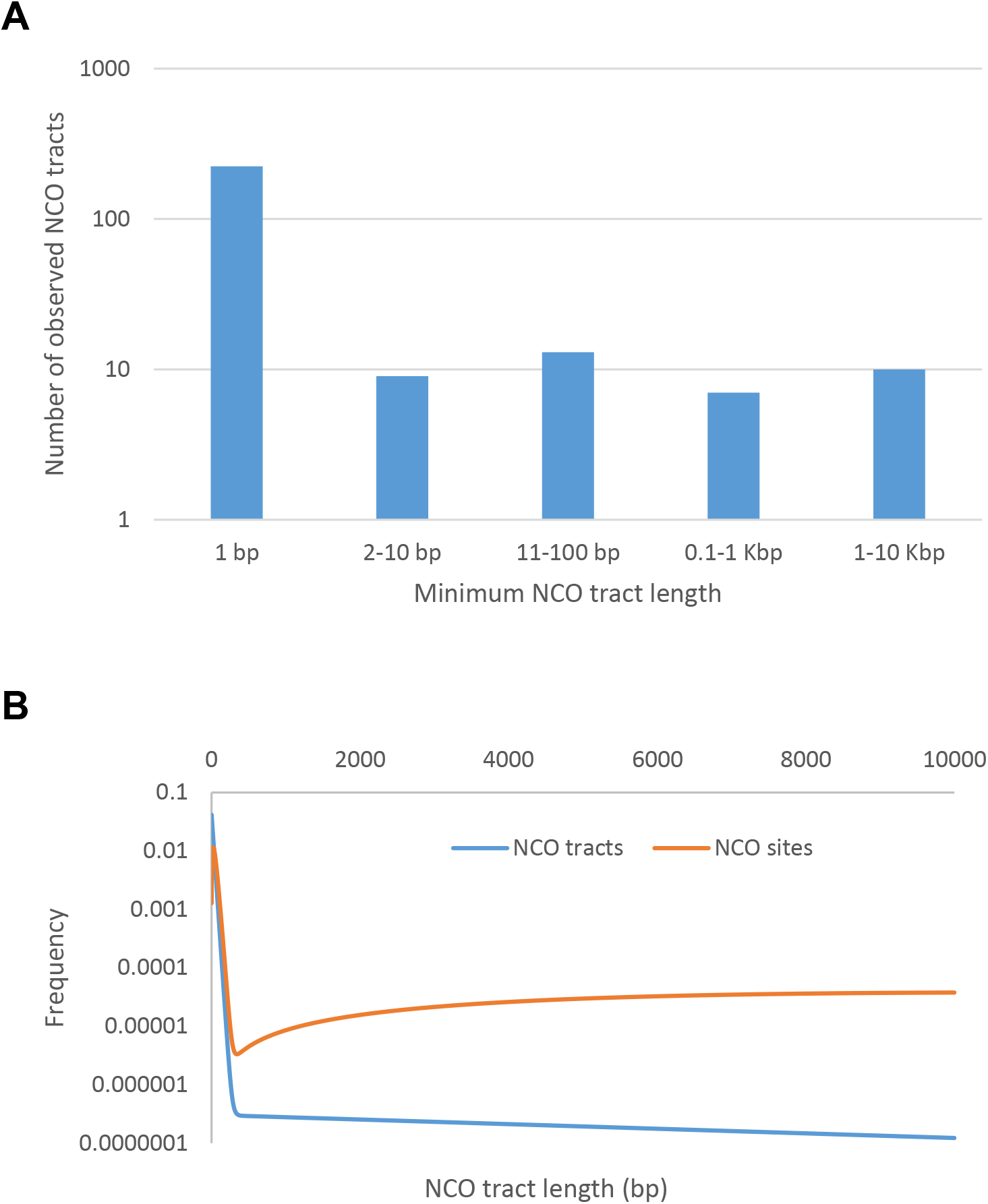
Distribution of NCO tract lengths, for actual data and best-fit model. (A) Minimum inferred lengths of observed NCO tracts shorter than 10 Kbp. (B) Distribution of NCO tract lengths for the best-fit mixture of two geometric distributions model (in blue), and weighted by tract length (in orange).

We next considered a more general scenario where NCO tract lengths are modeled as a mixture of two geometric distributions. This would be appropriate if NCO recombination could occur through two separate molecular pathways, each of which produced tracts whose length followed geometric distributions. Our maximum likelihood estimate had 99.8% of NCO tracts following a distribution with a mean length of 24 bp (95% CI = 18 – 31 bp), while the remaining NCO tracts had a mean length of 11 Kbp (95% CI = 3 – 100+ Kbp). Under this best-fit model, just 1.6% of NCO tracts are longer than 100 bp, but these longer tracts account for 31.6% of all sites that are converted by NCO recombination (see blue and orange curves in Figure 6B respectively).

## Discussion

Pedigree-based studies of recombination, while common in previous decades due to technological and computational limitations, have been mostly superseded now by studies that indirectly estimate recombination rates from patterns of linkage disequilibrium (e.g., Apuli et al. 2020; Beeson et al. 2019; Dreissig et al. 2019; Jones et al. 2019; Pfeifer 2020; Robinson et al. 2019; Schield et al 2020; Schwarzkopf et al. 2020; Shanfelter et al. 2019; Spence and Song 2019; Xue et al. 2020). We argue though that despite the substantial amount of time and effort required to conduct pedigree-based studies, they can provide invaluable information that is inaccessible by other methods. LD-based recombination estimates, by their nature, are averages across time and individuals, are influenced by any evolutionary force that affects patterns of genetic variation (e.g., changes in population size, migration, admixture, natural selection, etc.), and require assumptions about the effective population size to be converted into an actual per generation rate. They cannot provide any information on sex-specific differences in CO rates, nor are they very informative about NCO recombination. For example, while baboon pyrho estimates are, on average, slightly elevated in sub-telomeric regions, this obscures the observation that male crossovers are 10-15 times more prevalent than female crossovers in the distal 5 Mbp of each chromosome arm. Human pedigree studies show the same general pattern (Broman et al. 1998; Kong et al. 2002), but the sex-bias is much larger in baboons than in humans.

One ancillary benefit of our pedigree-based examination of recombination in baboons is that it helped provide some independent information on the quality of the existing Panubis1.0 genome assembly. Panubis1.0 utilized a combination of Illumina short-read, Oxford Nanopore long-read, 10x Genomics linked-read, Bionano optical map and Hi-C sequence data to create an assembly with N50 contig size of 1.46 Mbp and single scaffolds that span each of the autosomes (Batra et al. 2020). The Hi-C data in particular enabled Panubis1.0 to be a truly *de novo* genome assembly, unlike the previous reference-guided baboon assembly (Rogers et al. 2019). However, there is some concern that Hi-C based scaffolding is susceptible to incorrect orientation of contigs, leading to inversion errors in the resulting assembly (e.g., Burton et al. 2013). Here, traditional linkage analyses enabled us to identify more than 20 likely assembly errors, most of which were putative inversions (Supplementary Table S3). This suggests that caution should be taken in accepting Hi-C based scaffolding without the presence of orthogonal sources of corroborating data.

Evidence for an inversion assembly error in linkage data comes from the presence of three closely spaced crossovers in one or more individuals (Broman et al. 1998, 2003; Figure 2). Six of these cases are also consistent with a single crossover associated with a long (24 – 86 Kbp), nearby NCO tract. While the data that we have cannot rule out either of these explanations, the relative dearth of putative long NCO tracts that are not associated with potential genome assembly errors strongly suggests that most (if not all) of the apparent long baboon NCO tracts are artefacts and not real. Similarly, we postulate that at least some of the long human NCO tracts identified in previous studies (Williams et al. 2015; Halldorsson et al. 2016) are actually due to microassembly errors or polymorphic structural variants. It is also likely that some of them represent complex NCO events (with multiple smaller conversion tracts) that are misclassified due to low marker density.

Finally, we note that even after removing all apparent long (e.g., > 10 Kbp) NCO tracts, our data show that a simple geometric model of NCO tract lengths is inappropriate, at least for baboons. While the mixture-of-two-geometric-distributions model we considered is somewhat arbitrary, it captures the qualitative observation that most baboon NCO tracts are quite short, but a small minority can be much longer. The extent to which our findings reflect general patterns of NCO recombination is unclear at this time. To date, only two other mammalian species have been studied in depth. Our results are qualitatively similar to the findings of NCO studies in humans (Williams et al. 2015; Halldorsson et al. 2016), but long NCO tracts seem to be much rarer in studies of hybrid mice (Li et al. 2019; Gergelits et al. 2021). High-resolution studies in additional species will be needed to better understand how empirical patterns of recombination vary across species, and whether additional molecular models (e.g., a crossover pathway without interference) may be necessary to explain the inferred patterns of recombination in large pedigrees.

## Materials and Methods

### Samples, sequencing and variant calling

All samples for this study are putative olive baboons (*Papio anubis*) from the pedigreed baboon colony housed at the Southwest National Primate Research Center (SNPRC). We extracted DNA from blood or tissue samples and sent them to MedGenome, Inc. for sequencing (using standard protocols and libraries) on Illumina HiSeq 4000 and X machines. We generated novel whole-genome sequence data from 23 individuals, generated additional sequence data from several previously published baboon genomes, and combined these with data from previous studies (Robinson et al. 2019; Wu et al. 2020) to obtain a final data set that included 66 baboons with a median of 35.6X depth of coverage. These samples are included in two large pedigrees (Figure 1) as well as in a panel of 36 unrelated olive baboons. SRA accession numbers for all sequences used in this study are presented in Supplementary Table S4 and archived in NCBI BioProject PRJNA433868.

For each sample, we mapped all sequenced reads to the Panubis1.0 genome assembly (Batra et al. 2020) using BWA MEM (Li and Durbin 2009) before marking duplicate reads with Picard (https://broadinstitute.github.io/picard) and then genotyping with HaplotypeCaller from the Genome Analysis Toolkit (GATK; McKenna et al., 2010). We then produced a joint genotype call set with GATK GenotypeGVCFs before applying filters. Specifically, we excluded sites in soft-masked regions of the genome, which correspond to repetitive and low complexity regions identified with WindowMasker (Morgulis et al., 2006), plus variants that were not bi-allelic SNPs. We also excluded genotypes with genotype quality (GQ) score less than 30, and sites with excess heterozygosity (defined as sites where the number of heterozygotes more than 3.5 standard deviations above random mating expectations). For the last criterion, we are aware that the 66 samples are not all unrelated, but since relatedness (and population structure) generally leads to less heterozygosity than random mating expectations, our approach is conservative.

### Pedigree-based identification of crossover events

We utilized the single nucleotide variants in the call set described above to identify recombination events from the meioses involving 10173, 12242, 9841, 1X2816, and their offspring (20 paternal meioses and 17 maternal meioses in total). Note that each sub-pedigree had a minimum of 5 offspring.

For a target meiosis, we first filtered the data to include only ‘informative’ sites where the parentally transmitted allele could be directly inferred. For example, when trying to identify paternal recombination events in 16517, we require the sire’s (10173) genotype to be heterozygous and the dam’s (12242) genotype to be homozygous. That way, the maternally transmitted allele is known and the paternally transmitted allele must be the other allele in 16517’s genotype. While the parental genome is unphased, it is straightforward to infer haplotypic phase by examining the patterns of alleles transmitted by the parent, and to identify potential recombination events by switches in which haplotype is inherited in each of the offspring (Coop et al. 2008). We employed additional filters by requiring genotype calls in all of a pedigree’s offspring, and by removing the 5 informative sites nearest to the ends of each chromosome.

We started by identifying all switches in transmitted haplotype that could be parsimoniously explained by a single crossover (Supplementary Table S1). We then manually examined all intervals (i.e., regions between consecutive informative markers) where we inferred the occurrence of two crossovers. Four of these intervals were long (e.g., > 60 Kb), not near chromosome ends (i.e., at least 2 Mb away), and involved a sub-pedigree with at least 7 offspring (i.e., with parent 10173, 1X2816 or 1X4519). For these, the evidence is quite strong that there were in fact two crossovers (rather than a genome assembly error or >2 crossovers). An additional eight intervals are shorter (2 – 30 Kb) and/or involve smaller sub-pedigrees with only 5 offspring. We have labelled these as ‘provisional’ crossovers, listed them in the “Provisional_COs” sheet in Supplementary Table S1, and included them for estimating the total genetic map length.

### Pedigree-based identification of genome assembly errors

We identified several unusual patterns in our crossover data that are suggestive of either genome assembly errors or polymorphic chromosomal rearrangements (Figure 2). For example, three closely spaced crossovers in a single meiosis are extremely unlikely due to crossover interference (Muller 1916), but could easily arise through the combination of a single crossover plus an inversion (Broman et al. 1998, 2003). We hypothesized that potential breakpoints are likely to occur in-between assembled contigs, and classified these synteny breaks as putative inversions (three crossovers within 10 Mb in the same meiosis), misplaced contigs (two closely linked crossovers occurring at the same locations in multiple individuals), translocations (misplaced contigs where the correct genomic location could be inferred), or single breaks of synteny (Supplementary Table S2). The genomic location of contig breaks within hypothesized breakpoints are shown in parentheses in the second and third columns of Table S2.

### LD-based recombination estimates

We used pyrho (Spence and Song 2019) to estimate local recombination rates from patterns of linkage disequilibrium in a panel of 36 unrelated olive baboon founders. Recombination rate inference with pyrho requires a demographic model, and for this we used SMC++ (Terhorst et al., 2017). For this analysis, we included genotypes with a minimum quality score of 40 and read depth ≥8, and applied a series of filters based on recommendations from GATK to minimize the inclusion of errors (QUAL < 30.0, QD < 2.0, FS > 60.0, MQ < 40.0, MQRankSum < −12.5, ReadPosRankSum < −8.0, SOR > 3.0, ExcHet < 0.05), leaving 14.4 million variants in total. To run SMC++, we used a random set of ten individuals as the “distinguished” lineages, and set the polarization parameter (-p) to 0.5 to handle uncertainty in the polarization of derived versus ancestral alleles. We then followed the developer’s instructions to incorporate the demographic model from SMC++ into the recombination rate inference with pyrho. We also incorporated two additional filters before running pyrho; we excluded singletons, which are uninformative for LD-based recombination rate inference, and we thinned variants so that no two SNPs were closer than 10 bp, leaving 10.4 million variants in total. To handle the large number of haplotypes in our dataset (n=72), we enabled the Moran approximation with: --approx --moran_pop_size 98. After exploring a number of block penalty (smoothing) and window size parameters with hyperparam, we found that a block penalty of 10 and window size of 75 were optimal. After manual inspection of the output, we removed the pyrho value for a single interval on chromosome 19 (approximate positions 24.92 – 25.02 Mb) where the estimated genetic map length was ~102 cM. There is no evidence for any crossovers near this region, so we deemed the extremely large estimate to be unreliable.

pyrho and other LD-based methods most naturally estimate the population scaled recombination parameter ρ (= 4Nr, where N is the effective population size and r is the recombination rate per generation). So, conversion of these values to actual per generation recombination rate estimates requires assumptions about other fundamental parameters such as N and/or the mutation rate. With the assumptions described above, the pyrho estimated total genetic map length was several times shorter than the sex-averaged crossover-based genetic map length. To make these results more compatible with each other, we rescaled the pyrho values to have the same total autosomal map length (2,293 cM) as estimated from crossovers by multiplying all pyrho-based estimates by the constant 5.577. This preserves local patterns of recombination rate variability while acknowledging the large uncertainties in estimating past population sizes.

### Identifying non-crossover (NCO) recombination

NCO recombination can be identified from pedigree data in an analogous way as crossover identification. Specifically, single NCO tracts show up as two very tightly linked crossovers in a single individual, or equivalently as one or more closely linked sites where an offspring inherits one parental haplotype, surrounded on both sides by much larger regions where the offspring inherits the other parental haplotype. Finally, we arbitrarily fixed the maximum NCO tract length size as 10 Kb, and analyzed apparent larger NCO tracts separately (see Results).

Pedigree 1 contained three offspring (32043, 32849 and 33863) of the 2nd generation individuals used for estimating NCO rates (Figure 1). For all sites contained in putative NCO tracts involving the parents of these offspring (19181 and 19348), we checked for Mendelian inconsistencies across the whole pedigree as a limited way to test whether putative NCO tracts might be caused by sequencing/genotyping errors in the offspring. (We did not find any.) To reduce the effect of potential genotyping errors in the parent, we excluded sites with more than two segregating alleles, as well as all apparent NCO tracts that are shared across multiple half or full siblings.

### NCO tract length distribution

We use an approximate maximum-likelihood approach to estimating the probability of the data as a function of NCO tract length distribution parameters. Here the data consist of the pattern of which informative sites are converted (or not converted) for each autosome of each meiosis. We assume that both the probability of NCO recombination and the NCO tract length distribution do not change across base pairs or meioses, and fix the former at 7.52 * 10^-6^ per base pair per generation, as estimated below. Without loss of generality, we assume that NCO tracts are initiated at a certain base pair, and then continue along the chromosome 5’ to 3’ until they end.

Suppose D(Λ) is a specific NCO tract length distribution governed by parameter(s) Λ. If m(Λ) is the mean tract length given Λ, then the per base pair probability of initiation of an NCO tract of length k is

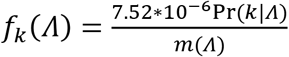

per meiosis. For a specific NCO tract, define {*o_k_*} as the number of k-mers (i.e., k consecutive base pairs) that overlap all of the informative sites converted in the tract and no others. Then the probability of observing an NCO tract is ∑*_k_f_k_* (*Λ*)*o_k_*. Similarly, define {*e_k_*} as the number of k-mers (across a chromosome) that must be excluded because they overlap with a non-converted site. The probability that none of the informative sites that shouldn’t be converted are actually converted is Π_*k*_(1 − *f_k_*(Λ))^*e_k_*^ ≈ *e*^−∑_*k*_*f_k_*(Λ)*e_k_*^. Finally, the likelihood of the full data is then the product of the separate likelihoods of observing each of the observed NCO tracts multiplied by the products of all of the probabilities of not converting the non-converted sites, across all meioses and autosomes.

Our analyses of actual data treated each contiguous NCO tract as separate, even if it was part of a complex NCO event, and arbitrarily required tract lengths to be 10 Kbp or shorter. This led to a total of 263 NCO tracts that we had information for. We first considered a geometric distribution for NCO tract lengths, requiring the mean length to be an integer. Confidence intervals were obtained using the standard asymptotic maximum likelihood assumptions. Next, we considered a mixture of two geometric distributions, parameterized by Λ = (α, m_1_, m_2_), where m_1_ and m_2_ are the means of the two distributions, and α is the proportion of tracts that have mean m_1_. We then calculated the likelihood of the data over a grid of parameter values, where α varied from 0 to 1 in increments of 0.001 and m_1_ and m_2_ varied from 10 – 100000 for all integers with 2 significant digits in this range. After obtaining the maximum likelihood estimate, we created profile likelihood curves for m_1_ and m_2_ to estimate approximate confidence intervals.

## Data Availability

All sequence data used in this study have been deposited in the Sequence Read Archive under NCBI BioProject PRJNA433868 with accession numbers given in Supplementary Table S4. The vcf file and pyrho genetic map will be made publicly available prior to publication.

## Acknowledgments

This work was supported by National Institutes of Health grants R24 OD017859 (to J.D.W. and L.A.C.) and R01 GM115433 (to J.D.W.).

## References

Apuli RP, Bernhardsson C, Schiffthaler B, Robinson KM, Jansson S, Street NR, Ingvarsson PK. 2020. Inferring the genomic landscape of recombination rate variation in European aspen (*Populus tremula*). G3 (Bethesda) 10:299–309.

Auton A, Fledel-Alon A, Pfeifer S, Venn O, Segurel L, Street T, Leffler EM, Bowden R, Aneas I, Broxholme J, et al. 2012. A fine-scale chimpanzee genetic map from population sequencing. Science 336:193–198.

Batra SS, Levy-Sakin M, Robinson J, Guillory J, Durinck S, Vilgalys TP, Kwok PY, Cox LA, Seshagiri S, Song YS, et al. 2020. Accurate assembly of the olive baboon (*Papio anubis*) using long-read and Hi-C data. Gigascience 9:giaa134.

Baudat F, de Massy B. 2007. Regulating double-stranded DNA break repair towards crossover or non-crossover during mammalian meiosis. Chromosome Res 15:565–577.

Beeson SK, Mickelson JR, McCue ME. 2019. Exploration of fine-scale recombination rate variation in the domestic horse. Genome Res 29:1744–1752.

Broman KW, Murray JC, Sheffield VC, White RL, Weber JL. 1998. Comprehensive human genetic maps: individual and sex-specific variation in recombination. Am J Hum Genet. 63:861–869.

Broman KW, Matsumoto N, Giglio S, Martin CL, Roseberry JA, Zuffardi O, Ledbetter DH, Weber JL. 2003. Coomon long inversion polymorphism on chromosome 8p. In: Goldstein DR, editor. Science and statistics: a festschrift for Terry Speed. IMS Lecture Notes-Monograph Series. pp. 237–245.

Burton JN, A Adey, Patwardhan RP, Qiu R, Kitzman JO, Shendure J. 2013. Chromosome-scale scaffolding of de novo genome assemblies based on chromatin interactions. Nat Biotechnol 12:1119–1125.

Chakravarti A, Buetow KH, Antonarakis SE, Waber PG, Boehm CD, Kazazian HH. 1984. Nonuniform recombination within the human beta-globin gene cluster. Am J Hum Genet 36:1239–1258.

Cole F, Keeney S, Jasin M. 2010. Comprehensive, fine-scale dissection of homologous recombination outcomes at a hot spot in mouse meiosis. Mol Cell 39:700–710.

Comeron JM, Ratnappan R, Bailin S. 2012. The many landscapes of recombination in *Drosophila melanogaster*. PLoS Genet 8:e1002905.

Coop G, Wen XQ, Ober C, Pritchard JK, Przeworski M. 2008. High-resolution mapping of crossovers reveals extensive variation in fine-scale recombination patterns among humans. Science 319:1395–1398.

Crawford DC, Bhangale T, Li N, Hellenthal G, Rieder MJ, Nickerson DA, Stephens M. 2004. Evidence for substantial fine-scale variation in recombination rates across the human genome. Nat Genet 36:700–706.

Dreissig S, Mascher M, Heckmann S. 2019. Variation in recombination rate is shaped by domestication and environmental conditions in barley. Mol Biol Evol 36:2029–2039.

Duret L, Galtier N. 2009. Biased gene conversion and the evolution of mammalian genomic landscapes. Annu Rev Genomics Hum Genet. 10:285–311.

Frisse L, Hudson RR, Bartoszewicz A, Wall JD, Donfack J, Di Rienzo A. 2001. Gene conversion and different population histories may explain the contrast between polymorphism and linkage disequilibrium levels. Am J Hum Genet 69:831–843.

Galtier N, Duret L. 2007. Adaptation or biased gene conversion? Extending the null hypothesis of molecular evolution. Trends Genet 23:273–277.

Gay J, Myers S, McVean G. 2007. Estimating meiotic gene conversion rates from population genetic data. Genetics 177:881–894.

Gergelits V, Parvanov E, Simecek P, Forejt J. 2021. Chromosome-wide characterization of meiotic noncrossovers (gene conversions) in mouse hybrids. Genetics 217:1–14.

Halldorsson BV, Hardarson MT, Kehr B, Styrkarsdottir U, Gylfason A, Thorleifsson G, Zink F, Jonasdottir Ad, Jonasdottir As, Sulem P, et al. 2016. The rate of meiotic gene conversion varies by sex and age. Nat Genet 48:1377–1384.

Hilliker AJ, Harauz G, Reaume AG, Gray M, Clark SH, Chovnick A. 1994. Meiotic gene conversion tract length distribution within the rosy locus of *Drosophila melanogaster*. Genetics 137:1019–1026.

Jasinska AJ, Service S, Levinson M, Slaten E, Lee O, Sobel E, Fairbanks LA, Bailey JN, Jorgensen MJ, Breidenthal SE, et al. 2007. A genetic linkage map of the vervet monkey (*Chlorocebus aethiops sabaeus*). Mamm Genome 18:347–360.

Jeffreys AJ, Kauppi L, Neumann R. 2001. Intensely punctate meiotic recombination in the class II region of the major histocompatibility complex. Nat Genet 29:217–222.

Jeffreys AJ, May CA. 2004. Intense and highly localized gene conversion activity in human meiotic crossover hot spots. Nat Genet 36:151–156.

Jeffreys AJ, Neumann R. 2005. Factors influencing recombination frequency and distribution in a human meiotic crossover hotspot. Hum Mol Genet 14:2277–2287.

Jones JC, Wallberg A, Christmas MJ, Kapheim KM, Webster MT. 2019. Extreme differences in recombination rate between the genomes of a solitary and a social bee. Mol Biol Evol 36:2277–2291.

Kong A, Gudbjartsson DF, Sainz J, Jonsdottir GM, Gudjonsson SA, Richardsson B. 2002. A high-resolution recombination map of the human genome. Nat Genet. 31:241–247.

Kong A, Barnard J, Gudbjartsson DF, Thorleifsson G, Jonsdottir G, Sigurdardottir S, Richardsson B, Jonsdottir D, Thorgeirsson T, Frigge ML, et al. 2004. Recombination rate and reproductive success in humans. Nat Genet 36:1203–1206.

Li H, Durbin R. 2009. Fast and accurate short read alignment with Burrows-Wheeler transform. Bioinformatics 25:1754–1760.

Li R, Bitoun E, Altemose N, Davies RW, Davies B, Myers SR. 2019. A high-resolution map of non-crossover events reveals impacts of genetic diversity on mammalian meiotic recombination. Nat Comm 10:3900.

Martin HC, Christ R, Hussin JG, O’Connell J, Gordon S, Mbarek H, Hottenga JJ, McAloney K, Willemsen G, Gasparini P, et al. 2015. Multicohort analysis of the maternal age effect on recombination. Nat Commun 6:7846.

McKenna A, Hanna M, Banks E, Sivachenko A, Cibulskis K, Kernytsky A, Garimella K, Altshuler D, Gabriel S, Daly M, et al. 2010. The Genome Analysis Toolkit: a MapReduce framework for analyzing next-generation DNA sequencing data. Genome Res 20:1297–1303.

Miller DE, Takeo S, Nandanan K, Paulson A, Gogol MM, Noll AC, Perera AG, Walton KN, Gilliland WD, Li H, et al. 2012. A whole-chromosome analysis of meiotic recombination in *Drosophila melanogaster*. G3 (Bethesda) 2:249–260.

Morgulis A, Gertz EM, Schäffer AA, Agarwala R. 2006. WindowMasker: window-based masker for sequenced genomes. Bioinformatics 22:134–141.

Muller HJ (1916) The mechanism of crossing over. Am Nat 50: 193–221.

Myers S, Bottolo L, Freeman C, McVean G, Donnelly P. 2005. A fine-scale map of recombination rates and hotspots across the human genome. Science 310:321–324.

Narasimhan VM, Rahbari R, Scally A, Wuster A, Mason D, Xue Y, Wright J, Trembath RC, Maher ER, van Heel DA, et al. 2017. Estimating the human mutation rate from autozygous segments reveals population differences in human mutational processes. Nat Commun 8:303.

Odenthal-Hesse L, Berg IL, Veselis A, Jeffreys AJ, May CA. 2014. Transmission distortion affecting human noncrossover but not crossover recombination: a hidden source of meiotic drive. PLoS Genet 10:e1004106.

Orr-Weaver TL, Szostak JW, Rothstein RJ. 1981. Yeast transformation: a model system for the study of recombination. Proc Natl Acad Sci USA 78:6354–6358.

Padhukasahasram B, Rannala B. 2013. Meiotic gene-conversion rate and tract length variation in the human genome. Eur J Hum Genet https://doi.org/10.1038/ejhg.2013.30.

Paigen K, Petkov PM. 2018. PRDM9 and its role in genetic recombination. Trends Genet. 34:291–300.

Pfeifer SP. 2020. A fine-scale genetic map for vervet monkeys. Mol Biol Evol 37:1855–1865.

Ptak SE, Hinds DA, Koehler K, Nickel B, Patil N, Ballinger DG, Przeworski M, Frazer KA, Pääbo S. 2005. Fine-scale recombination patterns differ between chimpanzees and humans. Nat Genet. 37:429–434.

Robinson JA, Belsare S, Birnbaum S, Newman DE, Chan J, Glenn JP, Ferguson B, Cox LA, Wall JD. 2019. Analysis of 100 high-coverage genomes from a pedigreed captive baboon colony. Genome Res 29:848–856.

Rogers J, Mahaney MC, Witte SM, Nair S, Newman D, Wedel S, Rodriguez LA, Rice KS, Perelygin A, Slifer M, et al. 2000. A genetic linkage map of the baboon (*Papio hamadryas*) genome based on human microsatellite polymorphisms. Genomics 67:237–247.

Rogers J, Garcia R, Shelledy W, Kaplan J, Arya A, Johnson Z, Bergstrom M, Novakowski L, Nair P, Vinson A, et al. 2006. An initial genetic linkage map of the rhesus macaque (*Macaca mulatta*) genome using human microsatellite loci. Genomics 87:30–38.

Rogers J, Raveendran M, Harris RA, Mailund T, Leppälä K, Athanasiadis G, Schierup MH, Cheng J, Munch K, Walker JA, et al. 2019. The comparative genomics and complex population history of *Papio* baboons. Sci Adv 5:eaau6947.

Schield DR, Pasquesi GIM, Perry BW, Adams RH, Nikolakis ZL, Westfall AK, Orton RW, Meik JM, Mackessy SP, Castoe TA. 2020. Snake recombination landscapes are concentrated in functional regions despite PRDM9. Mol Biol Evol 37:1272–1294.

Schwarzkopf EJ, Motamayor JC, Cornejo OE. 2020. Genetic differentiation and intrinsic genomic features explain variation in recombination hotspots among cocoa tree populations. BMC Genomics 21:332.

Shanfelter AF, Archambeault SL, White MA. 2019. Divergent fine-scale recombination landscapes between a freshwater and marine population of threespine stickleback fish. Genome Biol Evol 11:1573–1585.

Spence JP, Song YS. 2019. Inference and analysis of population-specific fine-scale recombination maps across 26 diverse human populations. Sci Adv 5:eaaw9206.

Stevison LS, Woerner AE, Kidd JM, Kelley JL, Veeramah KR, McManus KF, Great Ape Genome Project, Bustamante CD, Hammer MF, Wall JD. 2016. The time scale of recombination rate evolution in great apes. Mol Biol Evol. 33:928–945.

Szostak JW, Orr-Weaver TL, Rothstein RJ, Stahl FW. 1983. The double-strand break repair model for recombination. Cell 33:25–35.

Terhorst J, Kamm JA, Song YS. 2017. Robust and scalable inference of population history from hundreds of unphased whole genomes. Nat Genet 49:303–309.

Venn O, Turner I, Mathieson I, de Groot N, Bontrop R, McVean G. 2014. Nonhuman genetics. Strong male bias drives germline mutation in chimpanzees. Science 344:1272–1275.

Wall JD, Pritchard JK. 2003. Haplotype blocks and linkage disequilibrium in the human genome. Nat Rev Genet 4:587–597.

Webb AJ, Berg IL, Jeffreys A. 2008. Sperm cross-over activity in regions of the human genome showing extreme breakdown of marker association. Proc Natl Acad Sci USA 105:10471–10476.

Williams AL, Genovese G, Dyer T, Altemose N, Truax K, Jun G, Patterson N, Myers SR, Curran JE, Duggirala R, et al. 2015. Non-crossover gene conversions show strong GC bias and unexpected clustering in humans. eLife 4:e04637.

Wu FL, Strand AI, Cox LA, Ober C, Wall JD, Moorjani P, Przeworski M. 2020. A comparison of humans and baboons suggests germline mutation rates do not track cell divisions. PLoS Biol 18:e3000838.

Xue C, Rustagi N, Liu X, Raveendran M, Harris RA, Venkata MG, Rogers J, Yu F. 2019. Reduced meiotic recombination in rhesus macaques and the origin of the human recombination landscape. PLoS One 15:e0236285.

Yin J, Jordan MI, Song YS. 2009. Joint estimation of gene conversion rates and mean conversion tract lengths from population SNP data. Bioinformatics 25:i231–i239.

