## Supplemental Table S2 for "High-resolution estimates of crossover and noncrossover recombination from a captive baboon colony"

**Table S2.** List of potential genome assembly errors identified from patterns of crossovers in pedigrees

| Chr | Breakpoint 1[1] | Breakpoint 2[1] | Parent(s) | Offspring | Type | pyrho confirmation? |
|---|---|---|---|---|---|---|
| 4 | 128908402-128926502 (128926121) | 129083010-129133287 (129115227) | 9841 | 15444 | Inversion | Yes |
| 5 | 87107633-87174522 (87174124) | 87763274-87834276 (87832318) | 1X4519 | 9128 | Inversion | No |
| 6 | 19187190-19205515 (19204753) | 19418591-19427887 (19426981) | 10173 | 18385 | Inversion | Yes |
| 7 | 152346-183891 (162649) | 578048-619612 (553186) | 10173 | 15444 | Inversion | Inconclusive |
| 7 | 2108730-2110031 (2109398) | 2181650-2183146 (2182558) | >3 | >8 | Misplaced contig | Yes |
| 7 | 96278371-96282033 (96279505) | 96803605-96811375 (96805194) | 1X4519, 1X2816 | 6955, 7625 | Inversion | Yes |
| 7 | 100778767-100780278 | 105498863-105705880 (105499543) | 1X4519 | 10987 | Inversion | No |
| 7 | 114482280-114482897 | 115014663-115030265 (115020065) | 1X4519 | 10987 | Inversion | No |
| 7 | 154337477-154346856 (154340366) | 156816763-156827272 (156821679) | 1X2816, 9841 | 10987, 19348 | Inversion | No |
| 7 | 156818082-156827354 (156821780) | 157535325-157538265 | 10173 | 26988, 28246 | Misplaced contig | Yes |
| 8 | 43476352-43564391 (43550211) | 43635387-43639241 | 10173, 1X2816 | >4 | Misplaced contig | Yes |
| 11 | 39803972-39849254 (39808739) | 40005564-40036643 (40010327) | 1X2816 | 10489 | Inversion | Yes |
| 12 | 101260717-101277410 (101270326) | 106479800-106525157 (106481018) | 12242 | 16517, 17903, 28246 | Inversion | Yes |
| 13 | 7037503-7174125 | 7552177-7575804 | 1X2816 | 8307 | Inversion | No |

| | | | | | | |
|---|---|---|---|---|---|---|
| | (7059365) | (7555231) | | | | |
| 13 | 76945690-77038672 | 77379993-77385849 | 1X2816 | 16517 | Inversion | N/A |
| 13 | 103472993-103476718 (103476237) | 104948032-104950546 (104948753) | 1X2816 | 7311 | Inversion | No |
| 13 | 103793201-103794244 | | 10173 | 15444, 17199 | Synteny break | Yes |
| 16 | 52176980-52180999 (52179410) | 52308031-52316803 (52308689) | 10173, 1X2816 | >5 | Translocation | Yes[2] |
| 20 | 3112639-3115037 (3114115) | 4152939-4154479 (4153923) | 10173 | 17903, 28246 | Inversion | No |
| 20 | 9060715-9068752 | 9382803-9391671 | 10173 | 18385 | Inversion | N/A |
| 20 | 13003097-13047006 (13004575) | | 10173, 1X2816 | >5 | Synteny break | N/A |
| 20 | 37314503-37350014 | 37712955-37718236 | 10173 | 26988 | Inversion | N/A |

[1] Region where breakpoint can be localized, with coordinates of gap between contigs in parentheses when available
[2] Region is likely syntenic with the end of chromosome 16 (after approximate position 89.67 Mb)
