## Supplemental Figure S1 for "High-resolution estimates of crossover and noncrossover recombination from a captive baboon colony"

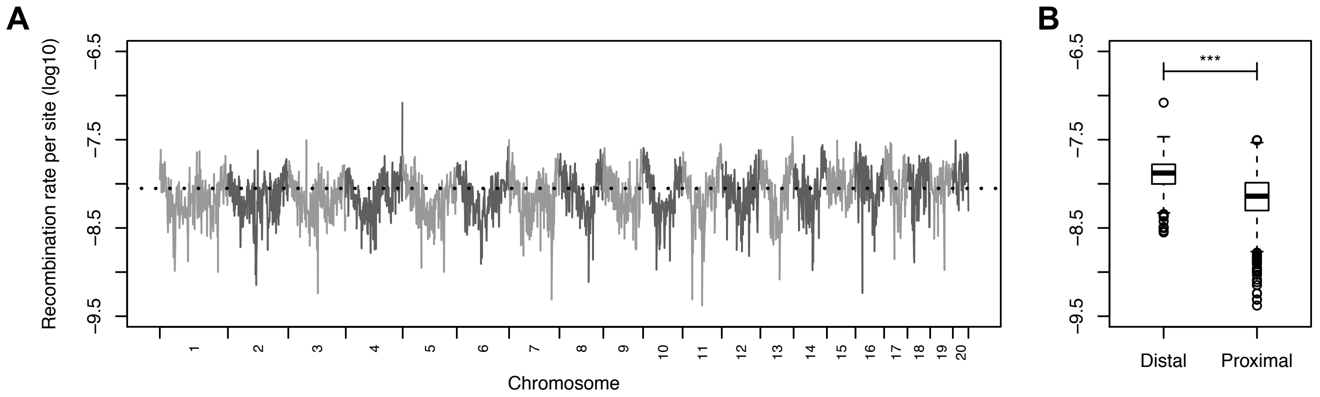


**Figure S1.** LD-based estimates of recombination using pyrho. (A) Average recombination rate estimates in non-overlapping 1 Mb windows across the genome. (B) Boxplots showing the significant difference in pyrho estimates between proximal (i.e., > 10 Mb from chromosome ends) and distal (<10 Mb from chromosome ends) windows.
